# Hearing through lip-reading: the brain synthesizes features of absent speech

**DOI:** 10.1101/395483

**Authors:** Mathieu Bourguignon, Martijn Baart, Efthymia C. Kapnoula, Nicola Molinaro

## Abstract

Lip-reading is crucial to understand speech in challenging conditions. Neuroimaging investigations have revealed that lip-reading activates auditory cortices in individuals covertly repeating absent—but known—speech. However, in real-life, one usually has no detailed information about the content of upcoming speech. Here we show that during silent lip-reading of *unknown* speech, activity in auditory cortices entrains more to absent speech than to seen lip movements at frequencies below 1 Hz. This entrainment to absent speech was characterized by a speech-to-brain delay of 50–100 ms as when actually listening to speech. We also observed entrainment to lip movements at the same low frequency in the right angular gyrus, an area involved in processing biological motion. These findings demonstrate that the brain can synthesize high-level features of absent unknown speech sounds from lip-reading that can facilitate the processing of the auditory input. Such a synthesis process may help explain well-documented bottom-up perceptual effects.

## Main text

In everyday situations, seeing the speaker’s articulatory mouth gestures, here referred to as lip-reading, can help us decode the speech signal.^1^ Human sensitivity to lip movements is so remarkable that even the auditory cortex seems to react to this visual input. In fact, seeing silent lip movements articulating simple speech sounds such as vowels or elementary words can activate the auditory cortex when participants are fully aware of what the absent auditory input should be.^2,3^ Thus, if prior knowledge about the absent sound is available, auditory information can be internally activated. However, in real-life, we usually have no detailed prior knowledge about what a speaker is going to say. Here, we tested the hypothesis that silent lip-reading provides sufficient information for the brain to synthesise relevant features of the absent speech.

When listening to natural continuous speech sounds, oscillatory cortical activity synchronises with the speech temporal structure in a remarkably faithful way.^4–8^ Such “speech entrainment” originates mainly from auditory cortices at frequencies matching phrasal (below 1 Hz) and syllable rates (4–8 Hz), and is thought to be essential for speech comprehension.^4,5,7,9^ Electroencephalography data have provided initial evidence that lip-reading might entrain cortical activity at syllable rate.^10^ However, it is unknown which brain areas were really involved in this process because Crosse and colleagues^10^ relied on the scalp distribution of the estimated entrainment. Most importantly, the reported syllable-level entrainment occurred only when detailed knowledge of the absent speech content was learned after multiple repetitions of the same stimulus. It is therefore unclear whether entrainment was driven by lip-reading, by covert production or repetition of the speech segment, by top-down lexical and semantic processes, or by a combination of these factors. Again, the critical question we addressed here is how the brain leverages lip reading to support speech processing in everyday situations where listeners *do not know* the content of the continuous speech signal.

We investigated entrainment to absent speech sounds when human adults were watching silent videos of a speaker telling an unknown story (see Fig. 1 for a comprehensive description of the experimental conditions). Brain signals were recorded by magnetoencephalography (MEG) and eye movements with an eye-tracker. Brain entrainment to both speech and lip-movements were quantified with coherence of MEG signals with speech temporal envelope and lip movements. Based on previous findings, we specifically expected to uncover speech entrainment at 0.5 Hz and 4–8 Hz,^4–8,11,12^ and lip entrainment at 2–5 Hz.^13,14^

**Figure 1.**
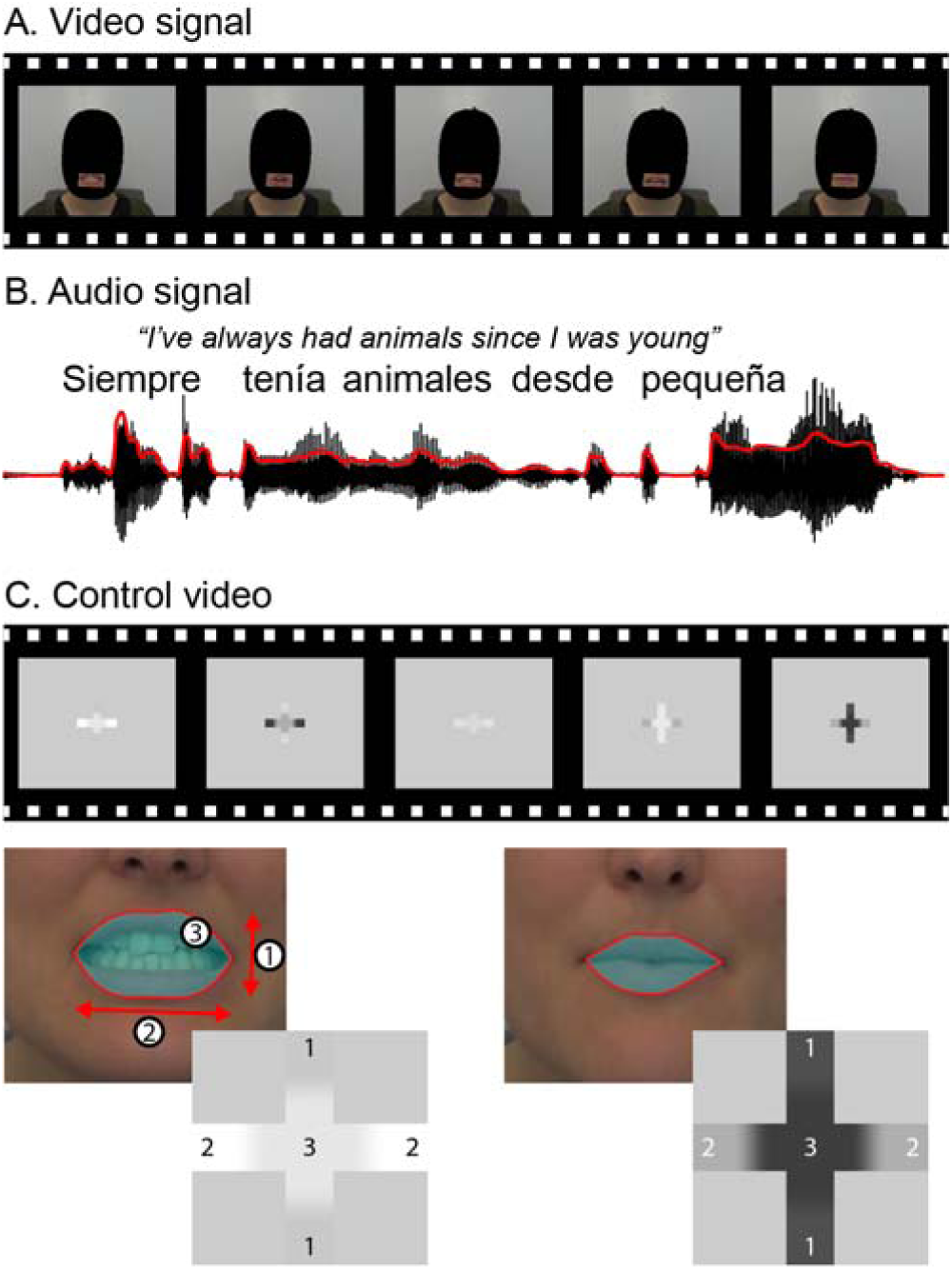
Experimental material. **A** and **B** — Two-second excerpt of video (**A**) and audio (**B**; speech temporal envelope in red) of the speaker telling a 5-min story about a given topic. There were 8 different videos. Video without sound was presented in *video-only*, and sound without video was presented in *audio-only*. **C —** Corresponding control video in which a flickering Greek cross encoded speaker’s mouth configuration. Based on a segmentation of mouth contours, the cross encoded mouth opening (1), mouth width (2), and mouth surface (3). The resulting video was presented in *control-video-only*.

## Results

In an *audio-only* condition, participants listened to a 5-min story while looking at a static fixation cross. Speech entrainment peaked above bilateral auditory regions at 0.5 Hz (Fig. 2A) and 4–8 Hz (Fig. 3), and underlying sources were located at bilateral auditory cortices for both 0.5 Hz (Fig. 2A and Table 1) and 4–8 Hz (Fig. 3 and Table 1).

**Table 1.** Significant peak of speech- and lip entrainment: peak MNI coordinates, significance level, confidence volume, and anatomical location. Only significant peaks of speech- (resp. lip-) entrainment that survived partialling out lip (resp. speech) are presented here. One exception is marked with * for which *p* = 0.063 after such correction.

| | Peak coordinates<br>[mm] | $p$ | Mean ± SD values | Confidence<br>volume [cm<sup>3</sup>] | Anatomical location |
| --- | --- | --- | --- | --- | --- |
| **Speech entrainment at 0.5 Hz** |  |  |  |  |  |
| Audio-only | [-64 -19 8] | $<10^{-3}$ | $0.076 \pm 0.045$ | 2.6 | Left auditory cortex |
| | [64 -21 6] | $<10^{-4}$ | $0.075 \pm 0.046$ | 5.5 | Right auditory cortex |
| Video-only | [-46 -30 11] | 0.003 | $0.025 \pm 0.017$ | 35.5 | Left auditory cortex |
| | [68 -14 -2] | 0.029 | $0.024 \pm 0.015$ | 5.6 | Right auditory cortex |
| | [-57 25 15] | 0.005 | $0.021 \pm 0.013$ | 9.6 | Left inferior frontal gyrus |
| | [-58 -15 41]* | 0.018 | $0.023 \pm 0.012$ | 21.3 | Left inferior precentral sulcus |
| **Lip entrainment at 0.5 Hz** |  |  |  |  |  |
| Video-only | [49 -46 10] | 0.002 | $0.022 \pm 0.014$ | 30.9 | Right angular gyrus |
| Control-video-only | [10 -89 -21] | $<10^{-3}$ | $0.028 \pm 0.023$ | 6.3 | Inferior occipital area |
| | [25 -96 -1] | 0.008 | $0.027 \pm 0.023$ | 11.7 | Right lateral occipital cortex |
| | [-23 -97 -4] | 0.046 | $0.023 \pm 0.014$ | 39.1 | Left lateral occipital cortex |
| **Speech entrainment at 4–8 Hz** |  |  |  |  |  |
| Audio-only | [-64 -18 7] | $<10^{-3}$ | $0.013 \pm 0.005$ | 1.4 | Left auditory cortex |
| | [67 -13 5] | $<10^{-3}$ | $0.020 \pm 0.009$ | 0.3 | Right auditory cortex |
| **Lip entrainment at 2–5 Hz** |  |  |  |  |  |
| Video-only | [-14 -97 11] | $<10^{-3}$ | $0.018 \pm 0.007$ | 8.3 | Left calcarine cortex |
| | [2 -93 -2] | $<10^{-3}$ | $0.018 \pm 0.008$ | 15.1 | Calcarine cortex |
| Control-video-only | [-1 -98 11] | $<10^{-3}$ | $0.026 \pm 0.016$ | 1.8 | Calcarine cortex |
| | [25 -97 -8] | $<10^{-3}$ | $0.025 \pm 0.012$ | 1.9 | Right lateral occipital cortex |
| | [-28 -94 -10] | $<10^{-3}$ | $0.024 \pm 0.012$ | 0.3 | Left lateral occipital cortex |

**Figure 2.**
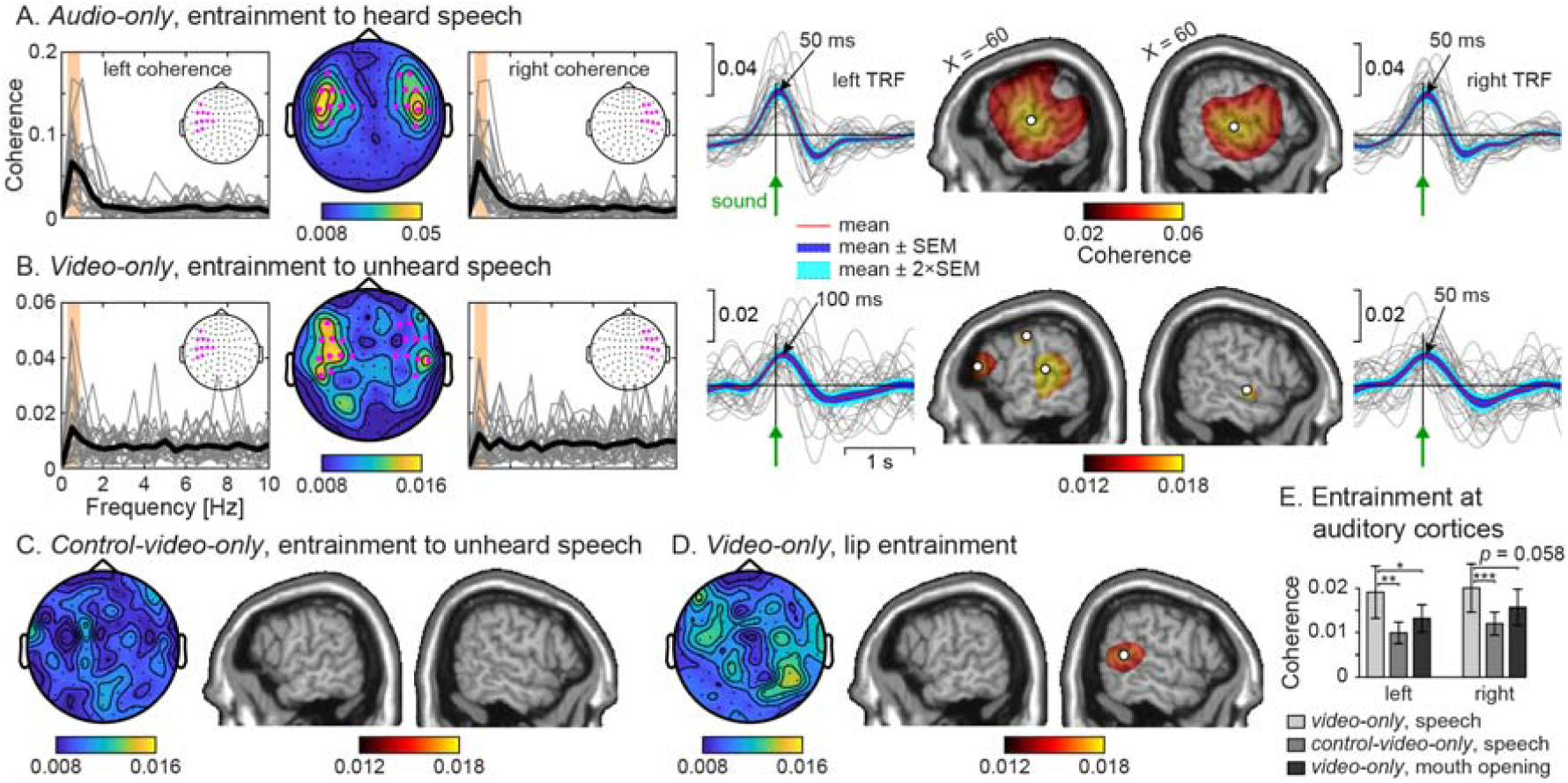
Speech and lip entrainment at 0.5 Hz. **A —** Speech entrainment in *audio-only*. The first three plots display the sensor distribution of speech entrainment at 0.5 Hz quantified with coherence and its spectral distribution at a selection of 10 left- and right-hemisphere sensors of maximal 0.5 Hz coherence (highlighted in magenta). Gray traces represent individual subjects’ spectra at the sensor of maximum 0.5 Hz coherence within the preselection, and the thick black trace is their group average. The last four plots display the brain distribution of significant speech entrainment quantified with coherence and the temporal response function (TRF) associated to speech temporal envelope at coordinates of peak coherence in the left and right hemispheres (marked with white discs). In brain maps, significant coherence values at MNI coordinates |X| > 40 mm were projected orthogonally onto the parasagittal slice of coordinates |X| = 60 mm. **B —** Same as in **A** for *video-only*, illustrating that seeing speaker’s face was enough to elicit significant speech entrainment at auditory cortices. Note that coherence spectra were estimated at the subject-specific sensor selected based on coherence in *audio-only* **C —** Sensor and source topography of speech entrainment in *control-video-only* wherein speech entrainment was not significant **D —** Sensor and source topography of lip entrainment in *video-only*. Lip entrainment was significant only in the right angular gyrus. **E** — Entrainment values at coordinates identified in *audio-only* (mean ± SD across participants).

**Figure 3.**
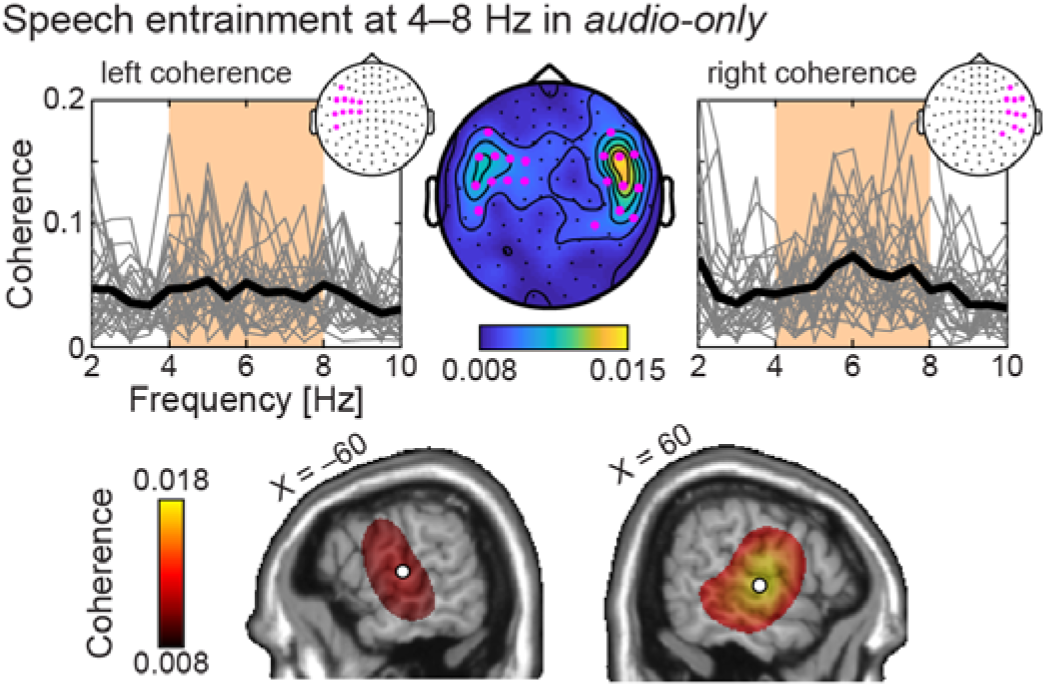
Speech entrainment at 4–8 Hz in *audio-only*. The top three plots display the sensor distribution of speech entrainment at 4–8 Hz quantified with coherence and its spectral distribution at a selection of 10 left- and right-hemisphere sensors of maximum coherence (highlighted in magenta). Gray traces represent individual subject’s spectra at the sensor of maximum 4–8 Hz coherence within the preselection, and the thick black trace is their group average. The two bottom plots display the brain distribution of significant speech entrainment. Significant coherence at MNI coordinates |X| > 40 mm were projected orthogonally onto the parasagittal slice of coordinates |X| = 60 mm. White discs are placed at the location of peak coherence in the left and right hemispheres.

In the *visual-only* condition, participants were looking at a silent video of the speaker telling a different 5-min story. Still, speech entrainment at 0.5 Hz occurred with the speech sound that was actually produced by the speaker, but that was not heard by the participants (Fig. 2B). The significant loci of 0.5-Hz speech entrainment were in bilateral auditory cortices, in the left inferior frontal gyrus, and in the inferior part of the left precentral sulcus (Fig. 2B and Table 1). Critically, the location of the auditory sources of maximum entrainment did not differ significantly between *audio-only* and *video-only* (left, *F*_3,998_ = 1.62, *p* = 0.18; right, *F*_3,998_ = 0.85, *p* = 0.47). Not surprisingly, the magnitude of 0.5-Hz speech entrainment was higher in *audio-only* than in *video-only* (left, *t*_27_ = 6.36, *p* < 0.0001; right, *t*_27_ = 6.07, *p* < 0.0001). Nevertheless, brain responses associated with speech entrainment at ∼0.5 Hz displayed a similar time-course, peaking with a delay of 50–100 ms with respect to speech sound in both *audio-only* and *video-only* (see Fig. 2A and 2B). These results demonstrate that within the auditory cortices, neuronal activity at ∼0.5 Hz is modulated similarly by heard speech sounds, and by absent speech sounds during lip-reading. Next, we addressed two decisive questions about this effect. 1) Is it specific to seeing speaker’s face or can it be triggered by any visual signal that features the temporal characteristics of lip movements? 2) Does it occur because speech temporal envelope is synthesised through a mapping from lip movements onto sound features, or alternatively, is it merely a direct consequence of lip-read induced visual activity that is fed forward to auditory areas?

Analysis of a *control-visual-only* condition revealed that entrainment to absent speech at auditory cortices was specific to seeing the speaker’s face. In the control condition, participants were looking at a silent video of a flickering Greek cross whose luminance pattern dynamically encoded speaker’s mouth configuration. We observed luminance-driven entrainment at 0.5 Hz at occipital cortices (Table 1), but no significant entrainment with absent speech (*p* > 0.1, Fig. 2C). Importantly, speech entrainment at auditory sources coordinates identified in *audio-only* was significantly higher in *video-only* than in *control-video-only* (left, *t*_27_ = 3.44, *p* = 0.0019; right, *t*_27_ = 4.44, *p* = 0.00014, see Fig. 2E). These differences in speech entrainment cannot be explained by differences in attention as participants did attend to the flickering cross in *control-video-only* about as much as speaker’s eyes and mouth in *video-only* (81.0 ± 20.9% vs. 87.5 ± 17.1%; *t*_26_ = 1.30, *p* = 0.20). This demonstrates that entrainment to absent speech in auditory cortices is specific to seeing the speaker’s face.

Although driven by lip-reading, auditory cortical activity at 0.5 Hz in *visual-only* entrained more to the absent speech sounds than to seen lip movements. Indeed, speech entrainment was stronger than lip entrainment at the left auditory source coordinates identified in *audio-only* (*t*_27_ = 2.52, *p* = 0.018, see Fig. 2E). The same trend was observed at the right auditory source (*t*_27_ = 1.98, *p* = 0.058, see Fig. 2E). However, at 0.5 Hz, lip movements entrained brain activity in the right angular gyrus (Fig. 2D and Table 1), a visual integration hub implicated in biological motion perception.^15,16^ Note that the dominant source of lip and speech entrainment were ∼4 cm apart (*F*_3,998_ = 4.68, *p* = 0.0030). Still, despite being distinct, their relative proximity might be the reason why speech entrainment was only marginally higher than lip entrainment at the right auditory cortex. Indeed, due to issues inherent to reconstructing brain signals based on extracranial signals, lip entrainment estimated at the auditory cortex was artificially enhanced by its genuine source ∼4 cm away. This leads us to conclude that entrainment in bilateral auditory cortices occurred with absent speech sound rather than with seen lip movements. As a final support to that claim, speech entrainment still peaked at bilateral auditory cortices after partialling out lip movements, less than 3 mm away from sources observed without partialling out lip movements. This pattern of results strongly supports the view that speech temporal envelope is synthesized through lip-reading.

Our data do not suggest the presence of the reciprocal effect of visual cortex entrainment to unseen lip movements in *audio-only*. Indeed, significant lip entrainment at 0.5 Hz occurred in auditory cortices only, and disappeared when partialling out entrainment to speech temporal envelope. No significant lip entrainment in this condition was uncovered at any of the other tested frequency ranges: 2–5 Hz and 4–8 Hz.

Lip entrainment at 2–5 Hz trivially occurred in occipital cortices in *video-only* and *control-video-only* (Fig. 4 and Table 1).

**Figure 4.**
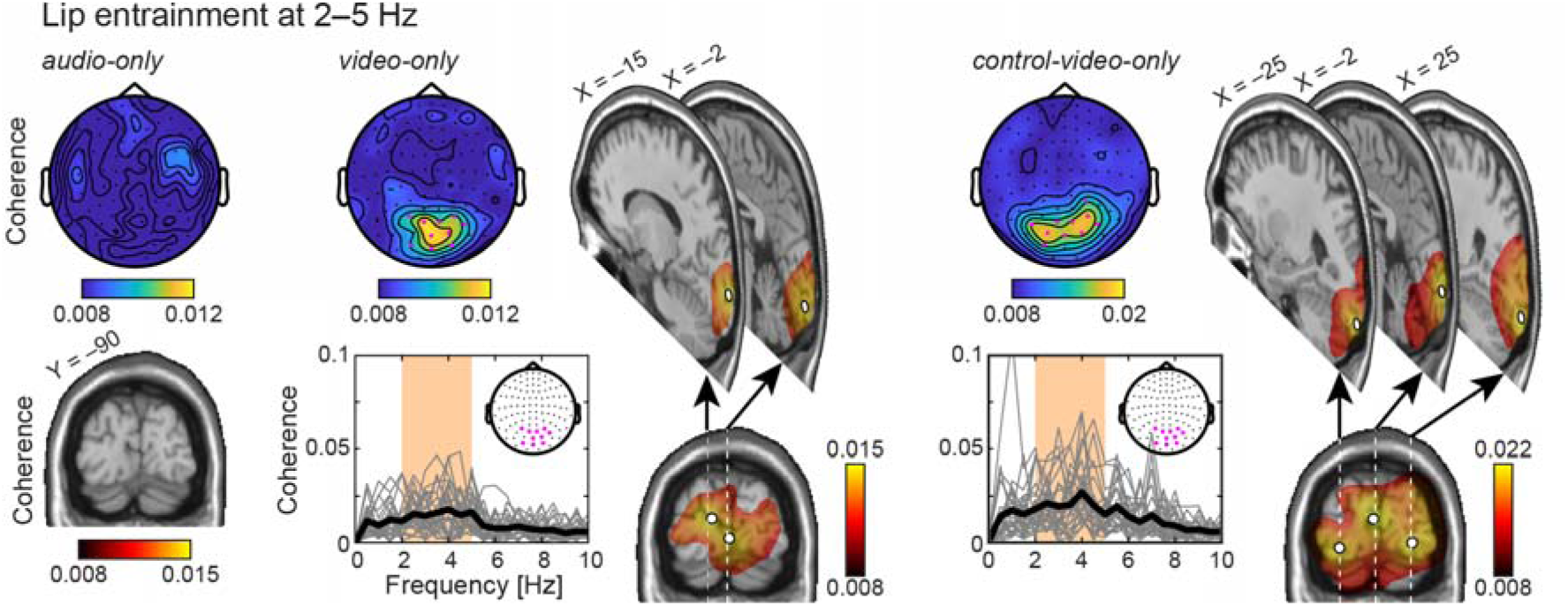
Lip entrainment at 2–5 Hz. Lip entrainment quantified with coherence is presented both in the sensor space and on the brain in all conditions (*audio-only, video-only, control-video-only*). In brain maps, significant coherence values at MNI coordinates Y < –70 mm were projected orthogonally onto the coronal slice of coordinates |Y| = 90 mm. Locations of peak coherence are marked with white discs. Note that coherence was not significant in *audio-only*. Additional parasagittal maps are presented for all significant peaks of coherence. In these maps, the orthogonal projection was performed for significant coherence values at Y coordinates less than 5 mm away from the selected slice Y coordinate. The figure also presents a spectral distribution of coherence at a selection of 10 sensors of maximum 2–5 Hz coherence (highlighted in magenta) in *video-only* and *control-video-only*. Gray traces represent individual subject’s spectra at the sensor of maximum 2–5 Hz coherence within the preselection, and the thick black trace is their group average.

At 4–8 Hz, there was significant speech entrainment in *video-only* and *control-video-only*, but only in occipital areas, and it vanished after partialling out the contribution of lip movements.

## Discussion

Altogether, the present data demonstrate that the brain can synthesise the slow (below 1 Hz) temporal dynamics of absent speech sounds from lip-reading. Watching silent articulatory mouth gestures without any prior knowledge about what the speaker is saying thus leaves a trace of the speech temporal envelope in auditory cortices that closely resembles that left by the actual speech sound.

### Entrainment to absent speech sounds in auditory cortices

The most striking finding of our study was that lip-reading induced entrainment in auditory cortices to the absent speech signal at frequencies below 1 Hz. Importantly, this entrainment was specific to lip-reading as it was not present in our visual-only control condition. Also, this entrainment was not a mere byproduct of entrainment to lip movements, as ruled out with partial coherence analysis. Instead, this genuine speech entrainment was strikingly similar to entrainment induced by actual speech: both are rooted in bilateral auditory cortices and are characterized by a speech-to-brain delay of 50–100 ms. This suggests the existence of a time-efficient synthesis mechanism based on which lip movements are mapped onto corresponding speech sound features. Such synthesis mechanism is likely grounded in the fact that the speech signal is coupled in—phonetic— space and time to the corresponding articulatory mouth gestures.^17–19^

Speech information in the frequency range below 1 Hz reflects the rhythmic alternation of intonational phrases. ^20^ Accordingly, corresponding entrainment to heard speech sounds has been hypothesised to reflect parsing or chunking of phrasal and sentential information.^7,8,21^ The most striking demonstration of this is that entrainment to phrasal and sentential rhythmicity is observed even for artificial stimuli in which phrasal and sentential prosody is absent.^21^ It has been argued that speech-brain coupling at frequencies below 1 Hz does not merely reflect entrainment to the acoustic properties of the external speech signal, but rather a process that organizes speech signal in higher-level linguistic structures.^22^ Interestingly, these brain mechanisms influence syntactic comprehension. For example, when listening to ambiguous sentences, 0–4 Hz auditory oscillations modulate the linguistic bias for grouping words into phrases.^23^ According to a recent view, low frequency speech entrainment would serve to align neural excitability with syntactic information to optimize language comprehension.^24^ If such speech-brain synchronization is not accurate, it can have a detrimental impact on the acquisition of typical language skills. In fact, the reduced phonological awareness in dyslexic readers has been associated to reduced speech entrainment at frequencies below 1 Hz.^25^ Overall, these findings highlight the fundamental role that this phrasal-level speech entrainment plays for language comprehension. In the present paper, we show that such high-level speech information can be retrieved based on lip-reading, and it is available for chunking and parsing lower-level linguistic units such has words, syllables and phonemes.

As 4–8-Hz frequencies match with syllable rate, corresponding speech entrainment has been hypothesised to reflect brain sensitivity to syllabic information. In support of this view, 4–8-Hz entrainment is enhanced when listening to intelligible speech compared to non intelligible speech.^4,5,9^ However, we did not observe such entrainment during silent lip-reading, which may suggest that the brain does not synthesise the detailed phonology of unfamiliar silent syllabic structures in that context. In support of this view, lip-reading is a very difficult task: even professional lip-readers struggle to decipher silent lip-read videos.^26^ This is presumably because different phonemes can be associated with very similar lip configurations (*e.g.*, /ba/, /pa/ and /ma/). However, when participants are primed (based on multiple repetitions) about the content of the audiovisual speech stimuli, this ambiguity in the mapping between articulatory mouth gestures and the corresponding phonemes disappears, and it has indeed been suggested that lip-reading can induce entrainment in auditory cortices at frequencies above 1 Hz in these conditions.^10^

The fact that we did not uncover lip-read-induced entrainement to absent speech sounds at syllable rate (4–8 Hz) does not imply that the brain does not synthesize fine-grained sound features in natural audiovisual conditions. Brain synthesis of fine-grained information might not surface in speech entrainment simply because temporal audiovisual coupling is highly variable: the delay between vision and sound varies by up to ∼100 ms.^27^ It is easy to show that such variability would conceal entrainment at 4–8 Hz while leaving essentially unaffected entrainment at frequencies below 1 Hz. Consequently, the below 1 Hz lip-read-induced entrainment to absent speech sounds could be the only visible trace of a synthesis process of the fine-grained features of absent sounds. If that speculation proves true, our results could help explain the McGurk illusion in which incongruent audio-visual information alters perceived sound identity.^28^ For example, listening to ‘ba’ while seeing somebody articulating ‘ga’ often leads to a fused percept of ‘da’. Through the time-efficient synthesis mechanism characterized by a delay of 50-100 ms between absent speech and brain oscillatory activity, seeing mouth movements could first activate the phonological representation of a subset of matching speech sounds. Then, upon processing of the speech sound, the brain would adopt a Bayesian strategy to select the most likely of them.^29^

### Entrainment to lip movements

During silent lip-reading, activity in early visual cortices entrained to lip movements mainly at 2–5 Hz. The same effect was previously identified in a study on audio-visual speech in which video and sound in each ear could be congruent or incongruent.^13^ The authors reported lip entrainment in early visual cortices that was modulated by audio-visual congruence, leading to the conclusion that occipital lip entrainment is modulated by attention.^13^ Lip entrainment in occipital cortices was also observed in a study that mainly focused on the neuronal basis of audio-visual speech perception in noise.^14^ This occipital entrainment dominant at 2–5 Hz is probably the first step necessary for the brain to synthesize features of the absent speech sounds. Our results suggest that corresponding signals are forwarded to the right angular gyrus.

The right angular gyrus was the dominant source of lip entrainment at frequencies below 1 Hz. This brain region is the convergence area for dorsal and ventral visual streams and is specialised for processing visual biological motion.^15^ It has been extensively shown to activate in response to observed hand movements in monkeys^30^ and in humans.^31^ It was also uncovered during observation of continuous goal-directed hand movements, based on an analysis analogous to that used in the present report: coherence between hand kinematics and brain signals recorded with MEG.^32^ Closer to our topic, the right angular gyrus activates during lip-reading^2,16,33^ and observation of mouth movements.^15^ Functionally, the angular gyrus would map visual input onto linguistic representation during reading,^34^ and also during lipreading.^35^ Our results shed light on the oscillatory dynamics underpinning such mapping during lip-reading: based on visual input at dominant lip movement frequencies (2–5 Hz), the angular gyrus presumably extracts below 1 Hz features of lip movements, which would serve as an intermediate step to synthesise speech sound features.

Previous studies looking into brain dynamics underlying lipreading of silent connected visual speech have essentially used speech known to the subjects, and focused on visuo-phonological mapping in occipital cortices.^35–37^ For example, a study quantified how well occipital 0.3–15-Hz EEG signals can be predicted based on motion changes, visual speech features, absent speech temporal envelope, and all possible combinations thereof.^36^ Overall, including absent speech temporal envelope improved such prediction. Also, visual activity has been reported to entrain more to absent speech sounds at 4–7 Hz when the video is played forward than backward.^35^ Importantly, that effect was not driven by entrainment to lip movements as lip entrainment was similar for videos played forward and backward. Instead, it was accompanied with increased top-down drive from left sensorimotor cortices to visual cortices. This was taken as evidence that visuo-phonological mapping occurs also in early visual cortices,^35,36^ through top-down mechanisms.^35^ However, in our data, lip-read-induced 4–8-Hz entrainment to absent speech in visual cortices disappeared after partialling out lip movements, indicating that it is weaker than entrainment to absent speech in auditory cortices at frequencies below 1 Hz. This is perhaps not surprising given that the primary signal for human speech is auditory, and not visual. Accordingly, cross-modal learning effects in speech are larger when lip-reading recalibrates the auditory perceptual system, than when a sound recalibrates the visual perceptual system.^38,39^ Our data shows that this asymmetry is also reflected in brain oscillatory activity.

### Conclusion

Our results shed light on the oscillatory dynamics underlying lip-reading. In line with previous studies,^13,14^ our data indicate that seeing lip movements first modulates neuronal activity in early visual cortices at their dominant frequencies (mainly in a 2–5 Hz range). Based on this activity, the right angular gyrus, which is the convergence area for dorsal and ventral visual streams and is specialised for processing visual biological motion,^15,16^ extracts slower features of lip movements. Finally, the slower dynamics in the lip movements are mapped onto corresponding speech sound features and this information is fed to auditory cortices. Receiving this information likely facilitates speech parsing, in line with the hypothesised role of entrainment to heard speech sounds at frequencies below 1 Hz.^7,8,21^ It might also explain the perceptual McGurk illusion in which lip-read information that is incongruent with the speech sound can alter perceived sound identity.^28^

## Methods

### Participants

Twenty-eight healthy adults native speakers of Spanish (17 females) aged 24.1±4.0 years (mean ± SD) were included in the study. All were right-handed according to self report, had normal or corrected to normal vision and normal hearing, had no prior history of neurological or psychiatric disorders, and were not taking any medication or substance that influenced the nervous system.

The experiment was approved by the BCBL Ethics Review Board and complied with the guidelines of the Helsinki Declaration. Written informed consent was obtained from all participants prior to testing.

### Experimental paradigm

Figure 1 presents stimulus examples and excerpts. The stimuli were derived from 8 audio-visual recordings of a native Spanish speaker talking for 5-min about a given topic (animals, books, food, holidays, movies, music, social media, and sports). Video and audio were simultaneously recorded with a digital camera (Canon Legria HF G10) and its internal microphone. Video recordings were framed as a head-shots, and recorded at a PAL standard of 25 frames per second (videos were 1920 × 1080 pixels in size, 24 bits/pixel, with an auditory sampling rate of 44100 Hz). The camera was placed ∼70 cm cm away from the speaker, and the face spanned about half of the vertical field of view. Final images were resized to a resolution of 1024 × 768 pixel.

For each video, a “control” video was created in which mouth movements were transduced into luminance changes (Fig. 1C). To that aim, we extracted lip contours from each individual frame of the video recordings with an in-house matlab code based on the approach of Eveno et al.^40^. In the “control” video, a Greek cross (300 × 300 pixels in size) changed its luminance according to mouth configuration (Fig. 1C). The center of the cross encoded mouth surface, its top and bottom portions encoded mouth opening, and its left- and rightmost portions encoded mouth width. In this configuration, the three represented parameters were spatially and temporally congruent with the portion of the mouth they parametrized. All portions were smoothly connected by buffers along which the weight of the encoded parameters varied as a squared cosine.

A “control” audio was also derived from initial sound recordings but related conditions are not reported here.

Participants underwent 10 experimental audio-visual conditions while sitting with their head in a MEG helmet. Nine of these conditions resulted from all possible combinations of 3 types of visual conditions (original, control, no-video) and 3 types of audio conditions (original, control, no-audio). The no-audio and no-video condition was trivially labeled as the *rest* condition. The other 8 conditions were assigned to the 8 stories so that the same story was never presented twice, and the condition–story assignment was counterbalanced across subjects. The tenth condition was a localizer condition in which participants attended 400-Hz pure tones and checkerboard pattern reversals. All conditions were presented in a random order. Videos were projected onto a back-projection screen (videos were 41 cm × 35 cm in size) placed in front of the participants at a distance of ∼1 m. Sounds were delivered at 60 dB (measured at ear-level) through a front-facing speaker (Panphonics Oy, Espoo, Finland) placed ∼1 m behind the screen. Participants’ were instructed to watch the videos attentively, and listen to the sounds.

To investigate our research hypotheses, we focussed on the following conditions: 1) the original audio with no video, referred to as *audio-only*, 2) the original video with no audio, referred to as *video-only*, 3) the control video with no audio, referred to as the *control-video-only*, and 4) the *rest*.

### Data acquisition

Neuromagnetic signals were acquired with a whole-scalp-covering neuromagnetometer (Vectorview; Elekta Oy, Helsinki, Finland) in a magnetically shielded room. The recording pass-band was 0.1–330 Hz and the signals were sampled at 1 kHz. The head position inside the MEG helmet was continuously monitored by feeding current to 4 head-tracking coils located on the scalp. Head position indicator coils, three anatomical fiducials, and at least 150 head-surface points (covering the whole scalp and the nose surface) were localized in a common coordinate system using an electromagnetic tracker (Fastrak, Polhemus, Colchester, VT, USA).

Eye movements were tracked with an MEG-compatible eye tracker (EyeLink 1000 Plus, SR Research). Participants were calibrated using the standard 9-point display and monocular eye movements were recorded at a sampling rate of 1 kHz. Eye-movements were recorded throughout the duration of all experimental conditions.

High-resolution 3D-T1 cerebral magnetic resonance images (MRI) were acquired on a 3 Tesla MRI scan (Siemens Medical System, Erlangen, Germany).

### Data analysis

#### MEG preprocessing

Continuous MEG data were first preprocessed off-line using the temporal signal space separation method (correlation coefficient, 0.9; segment length, 10 s) to suppress external interferences and to correct for head movements.^41,42^ To further suppress heartbeat, eye-blink, and eye-movement artifacts, 30 independent components^43,44^ were evaluated from the MEG data low-pass filtered at 25 Hz using FastICA algorithm (dimension reduction, 30; non-linearity, tanh). Independent components corresponding to such artifacts were identified based on their topography and time course and further projected out from MEG signals.

#### Coherence analysis

Coherence was estimated between MEG signals, and 1) speech temporal envelope, 2) mouth opening, 3) mouth width, and 4) mouth surface. Speech temporal envelope was obtained as the rectified sound signals low-pass filtered at 50 Hz, further resampled to 1000 Hz time-locked to the MEG signals (Fig. 1B). Continuous data from each condition were split into 2-s epochs with 1.6-s epoch overlap. We used overlapping epochs as that leads to decreased noise on coherence estimates.^45^ MEG epochs exceeding 5 pT (magnetometers) or 1 pT/cm (gradiometers) were excluded from further analyses to avoid contamination of our data by any other source of artifact that would not have been dealt with by the temporal signal space separation or independent component analysis based artifact suppression. These steps led to a number of artifact-free epochs of 732 ± 36 (mean ± SD across participants and conditions), and a one-way repeated measures ANOVA revealed no difference in this amount between conditions (*F*_2,54_ = 1.07, *p* = 0.35). Coherence was then estimated at the sensor level using the formulation of Halliday et al.^46^. Data from gradiometer pairs were combined in the direction of maximum coherence as done in Bourguignon et al.^47^. Using a similar approach, previous studies^4–7,11,12,48^ demonstrated significant coupling with speech temporal envelope at frequencies corresponding to the production rate of phrases (0.5 Hz) and syllable (4–8 Hz). Lip movements rather entrain the visual cortices at 2–5 Hz.^13,14^ We therefore focused on these specific frequency ranges by averaging coherence across the frequency bins they encompass. Coherence maps were also averaged across participants.

Note that we here report only on coherence estimated between MEG signals and 1) speech temporal envelope and 2) mouth opening. Although tightly related, these two later signals displayed a moderate degree of coupling, that peaked at 0.5 Hz, and 4–8 Hz (Fig. 5, left panel). Mouth opening and mouth surface were coherent at > 0.7 across the 0–10 Hz range (Fig. 5, middle panel) and yielded similar results. Mouth opening and mouth width displayed a moderate level of coherence (Fig. 5, right panel). Mouth width was nevertheless disregarded because it led to lower entrainment values than mouth opening.

**Figure 5.**
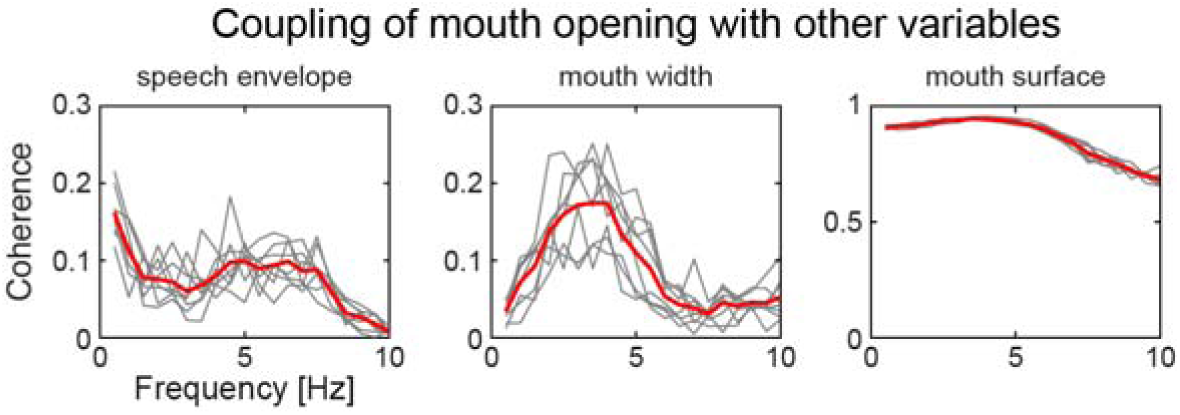
Frequency-dependent coupling of mouth opening with speech temporal envelope (*left*), mouth width (*middle*), and mouth surface (right). Coupling was quantified with coherence. There is one gray trace per video (8 in total), and thick red traces are the average across them all.

Coherence was also estimated at the source level. To do so, individual MRIs were first segmented using the Freesurfer software.^49^ Then, the MEG forward model was computed for two orthogonal tangential current dipoles placed on a homogeneous 5-mm grid source space covering the whole brain (MNE suite^50^). Coherence maps were produced within the computed source space at 0.5 Hz, 2–5 Hz, and 4–8 Hz using a linearly constrained minimum variance beamformer built based on *rest* data covariance matrix.^51,52^ Again, coherence in these two last ranges was obtained by simply averaging the coherence values across all frequency bins falling into these ranges. Source maps were then interpolated to a 1-mm homogenous grid and smoothed with a Gaussian kernel of 5-mm full-width-at-half-maximum. Both planar gradiometers and magnetometers were used for inverse modeling after dividing each sensor signal (and the corresponding forward-model coefficients) by its noise standard deviation. The noise variance was estimated from the continuous *rest* MEG data band-passed through 1–195 Hz, for each sensor separately.

Coherence maps were also produced at the group level. A non-linear transformation from individual MRIs to the MNI brain was first computed using the spatial normalization algorithm implemented in Statistical Parametric Mapping (SPM8^53,54^) and then applied to individual MRIs and coherence maps. This procedure generated a normalized coherence map in the MNI space for each subject and frequency range. Coherence maps were finally averaged across participants.

Individual and group-level coherence maps with speech temporal envelope (resp. mouth opening) were also estimated after controlling for mouth opening (resp. Speech temporal envelope). Partial coherence was used to this effect.^46^ Partial coherence is the direct generalization of partial correlation^55^ to the frequency domain.^46^

#### Estimation of temporal response functions

We used temporal response functions (TRFs) to model how speech temporal envelope affects the temporal dynamics of ∼0.5 Hz auditory cortical activity. A similar approach has been used to model brain responses to speech temporal envelope at 1–8 Hz,^56,57^ and to model brain responses to natural force fluctuations occurring during maintenance of constant hand grip contraction.^58^ TRFs are the direct analogue of evoked responses in the context of continuous stimulation.

We used the mTRF toolbox^59^ to estimate the TRF of auditory cortical activity associated with speech temporal envelope. The source signals considered were those at individual coordinates of maximum 0.5-Hz speech entrainment in audio-only, and in the orientation of maximum correlation between condition-specific source and sound envelope filtered through 0.2–1.5 Hz. Before TRF estimation, source signals were filtered through 0.2–1.5 Hz, sound envelope was convolved with a 50-ms square smoothing kernel, and both were down-sampled to 20 Hz. For each subject, the TRFs were modeled from –1.5 s to +2.5 s, for a fixed set of ridge values (λ = 2^0^, 2^1^, 2^2^… 2^20^). We adopted the following 10-fold cross-validation procedure to determine the optimal ridge value: For each subject, TRFs were estimated based on 90% of the data. TRFs were then multiplied by a window function with squared-sine transition on the edges (rising from 0 at –1.5 s to 1 at 1 s and ebbing from 1 at 2 s to 0 at 2.5 s) to dampen regression artifacts. We next used these windowed TRFs to predict the 10% of data left out, and estimated the Pearson correlation coefficient between predicted and measured signals. The square of the mean correlation value across the 10 runs provided an estimate of the proportion of variance explained by entrainment to speech temporal envelope. In the final analysis, we used the ridge value maximizing the mean explained variance across our 28 participants and both hemispheres in audio-only (λ = 2^13^). All TRFs were recomputed for this ridge value, and based on all the available data.

#### Eye tracking data

As in previous studies using eye tracking,^60,61^ eye-movements were automatically parsed into saccades and fixations using default psychophysical parameters. Adjacent saccades and fixations were combined into a single “look” that started at the onset of the saccade and ended at the offset of the fixation.

A region of interest was identified for each of the three critical objects: mouth and eyes in *video-only* and flickering cross in *control-video-only* (Fig. 6). In converting the coordinates of each look to the object being fixated, the boundaries of the regions of interest were extended by 50 pixels in order to account for noise and/or head-drift in the eye-track record. This did not result in any overlap between the eyes and mouth regions.

**Figure 6.**
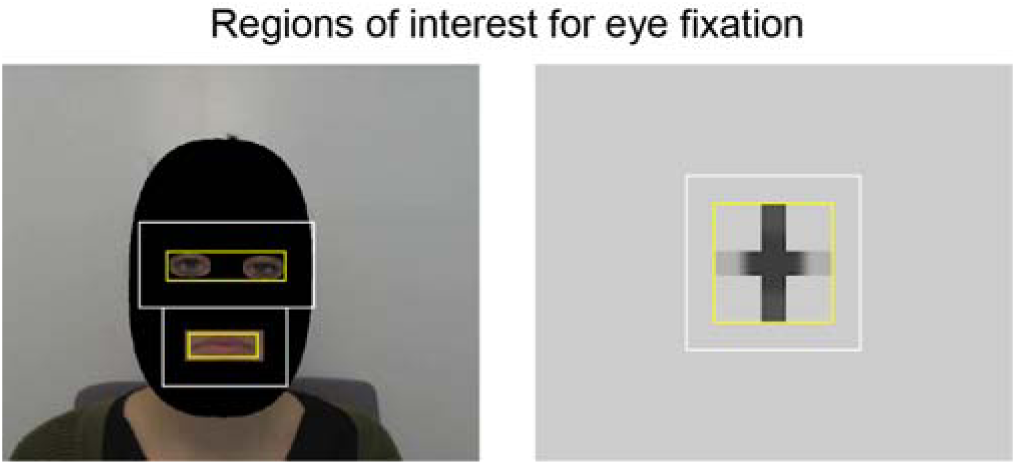
Regions of interest for eye fixation. The initial regions of interest are delineated in yellow, and the extended ones in white. Eye fixation analyses were based on extended regions. In *video-only* (*left*), the final region of interest comprised the mouth and the eyes. In control-video-only (*right*), it encompassed the flickering cross.

Based on these regions of interest, we estimated the proportion of eye fixation to the combined regions of interest encompassing eyes and mouth in *video-only* and flickering cross in *control-video-only*. Data from one participant were excluded due to technical issues during acquisition, and eye fixation analyses were thus based on data from 27 participants.

### Statistical analyses

The statistical significance of the local coherence maxima observed in group-level maps was assessed with a non-parametric permutation test that intrinsically corrects for multiple spatial comparisons.^62^ Subject- and group-level *rest* coherence maps were computed as done for the *genuine* maps, but with MEG signals replaced by *rest* MEG signals and sound or mouth signals unchanged. Group-level difference maps were obtained by subtracting *genuine* and *rest* group-level coherence maps. Under the null hypothesis that coherence maps are the same irrespective of the experimental condition, labeling *genuine* and *rest* are exchangeable at the subject-level prior to group-level difference map computation.^62^ To reject this hypothesis and to compute a threshold of statistical significance for the correctly labeled difference map, the permutation distribution of the maximum of the difference map’s absolute value was computed for a subset of 1000 permutations, for each hemisphere separately. The threshold at *p* < 0.05 was computed for each hemisphere separately as the 95^th^ percentile of the permutation distribution.^62^ All supra-threshold local coherence maxima were interpreted as indicative of brain regions showing statistically significant coupling with the audio or mouth signal.

A confidence volume was estimated for all significant local maxima, using the bootstrap-based method described in Bourguignon et al.^63^. The location of the maxima was also compared between conditions using the same bootstrap framework.^63^

For each local maximum, individual maximum coherence values were extracted within a 10-mm sphere centered on the group level coordinates, or on the coordinates of maxia in audio-only. Coherence values were compared between conditions or signals of reference with two-sided paired *t*-tests.

Individual proportions of fixations were transformed using the empirical-logit transformation.^64^ The fixations to eyes and mouth in *video-only* were compared to the fixations to the flickering cross in *control-video-only* using two-sided paired *t*-test across participants.

## Acknowledgment

Mathieu Bourguignon has been supported by the program Attract of Innoviris (grant 2015-BB2B-10), by the Spanish Ministry of Economy and Competitiveness (grant PSI2016-77175-P), and by the Marie Skłodowska-Curie Action of the European Commission (grant 743562). Martijn Baart has been supported by the Netherlands organization for scientific research (NWO, VENI grant 275-89-027). Efthymia C. Kapnoula has been supported by the Spanish Ministry of Economy and Competitiveness, through the Juan de la Cierva-Formación fellowship, and by the Spanish Ministry of Economy and Competitiveness (grant PSI2017-82563-P). Nicola Molinaro has been supported by the Spanish Ministry of Economy and Competitiveness (grant PSI2015-65694-P), the Agencia Estatal de Investigación (AEI), the Fondo Europeo de Desarrollo Regional (FEDER) and by the Basque government (grant PI_2016_1_0014). The authors acknowledge financial support from the Spanish Ministry of Economy and Competitiveness, through the “Severo Ochoa” Programme for Centres/Units of Excellence in R&D” (SEV-2015-490).

